# Egg hatching in the cestode *Schistocephalus solidus* shows no evidence of quorum sensing

**DOI:** 10.1101/2025.07.02.662852

**Authors:** Emily V. Kerns, Sara Engel, Panna A. Codner, Jesse N. Weber

**Affiliations:** Department of Integrative Biology, University of Wisconsin-Madison, Madison, Wisconsin, USA

## Abstract

*Schistocephalus solidus* is a parasitic cestode with a complex, multi-host life cycle. *S. solidus* reproduces in its terminal host either by exchanging gametes with similarly sized individuals or selfing. Fertilized eggs then pass through the feces of the host and hatch at the bottom of freshwater lakes. Previous work found that selfing greatly depresses egg hatching rates, presumably as a result of inbreeding depression. We predicted that *S. solidus* may have evolved quorum sensing (QS) during hatching as a mechanism to facilitate synchronized infection, thereby increasing the opportunity for outcrossing in its terminal host. We tested whether density-dependent hatch rates were present across three parasite populations, examining both outcrossed and selfed progeny. We predicted that if QS was present, it would be common across all populations, and that increasing egg density would result in higher hatching rates. We also expected that outcrossed eggs would hatch at higher rates than those produced via selfing. While we found different hatching rates between populations, there was no effect of egg density. Selfed eggs did hatch at significantly lower rates than outcrossed eggs, replicating previous findings. Although we failed to find density dependent hatching in our limited sample, we conclude by discussing the conditions in which QS may evolve in isolated *S. solidus* populations.

## Introduction

Despite their ubiquity and diversity, parasites are a historically understudied group of organisms. Although most research focuses on host biology, parasites are a strong force of natural selection that can drive drastic demographic shifts, and sometimes evolution, in hosts over ecological time scales.^1–3^ A long-standing model system for studying the evolution and ecology of parasitism is the threespine stickleback (*Gasterosteus aculeatus*) fish and its parasitic cestode *Schistocephalus solidus*.^4^ While *S. solidus* clearly imposes strong selection on stickleback to mount various forms of resistance and tolerance^5^, the cestode has received much less research attention than its fish host, particularly during its early life stages.

*S. solidus* was the first parasite observed to display a complex, trophically transmitted life cycle.^4,6^ Initially, the cestode’s eggs develop in freshwater before hatching into coracidia.^7^ The coracidia are ingested by its first intermediate host, copepods in the order Cyclopoida, and then migrate into the host body cavity before developing into procercoid larvae. For the parasite’s life cycle to progress, infected copepods must be ingested by threespine stickleback, the specific host for *S. solidus*. The majority of the cestode’s growth occurs within the stickleback as it develops into a plerocercoid larva.^4^ The infected fish is then predated on by a bird, *S. solidus’*s terminal host, where the parasite does not siphon any resources but rather immediately develops its reproductive anatomy and generates offspring in the gut via either cross fertilization or selfing.

The fertilized eggs are expelled via bird feces.^8,9^ As noted previously, while *S. solidus* can infect many species of copepods and birds, it can only successfully complete its life cycle if ingested by threespine stickleback, making this cestode unique in that its obligate host (threespine stickleback) is not its definitive host. However, because of the bird’s ability to move large distances while carrying the parasite, there is often gene flow across large geographic areas, supported by genetic evidence that *S. solidus* exhibit isolation-by-distance.^10^

As the potential for *S. solidus* to outcross is dependent on a bird ingesting multiple mature parasites, synchronized infection during its early life stages may be advantageous for the cestode to prevent inbreeding depression. Indeed, coinfections of threespine stickleback are common, but they also cause the cestodes to be smaller on average than individuals in singly infected fish, which reduces fecundity.^10,11^ Analogous to the effect of crowding in stickleback, procercoids in multiply infected copepods tended to be smaller and exhibited a slower growth rate. However, coracida were more likely to successfully establish infection in a copepod when there was a higher rate of *S. solidus* exposures.^12^ Additionally, inbreeding depression is extremely costly for *S. solidus*, with progeny of selfed individuals exhibiting less than 10% of the hatching rate of their outcrossed counterparts.^13^ Increased and synchronized infection early in the *S. solidus* life cycle may therefore facilitate coinfection in terminal hosts and increase the potential for outcrossing.

One mechanism to facilitate this beneficial infection synchrony might be the use of quorum sensing (QS) by *S. solidus* eggs. First described in bacterial cells,^14^ QS involves sensing local population density and then releasing a signal when a critical density is reached. While typically associated with bacteria, eukaryotic parasitic microbes, such as *Trypanosoma brucei*, have been shown to exhibit QS properties.^15^ In the case of *T. brucei*, QS signals induce a pleomorphic response where the parasites develop stumpy, non-dividing body types in its mammalian host. Given the cost of inbreeding depression and the presence of QS in other eukaryotic parasites, QS may have evolved in *S. solidus* to coordinate hatching.

While coordinating coracidia hatching may increase the potential for successful coinfection in a copepod, if and how *S. solidus* does this remains an open question. We therefore tested whether egg density influences hatching rates of *S. solidus* coracidia, predicting that eggs in higher densities would also exhibit higher hatch rates, presumably mediated by chemical signals emitted by hatched worms (i.e., a form of QS). We tested this hypothesis using laboratory-bred cestodes originating from three geographically distinct populations, which allowed us to assess whether QS in *S. solidus* is a common trait or is specific to some populations.

## Materials & Methods

### Egg Collection

We followed a previously described *S. solidus* breeding protocol.^16^ Briefly, mature cestodes were collected from infected fish in Walby Lake, Alaska, USA (61.617963, - 149.213644), Myvatn Lake, Iceland (65.596929, -17.002945), and Echo Lake, British Columbia, Canada (49.988258, -125.410090). Cestodes of similar size, which increases the likelihood of outcrossing^23^, were placed in sterile, nylon biopsy bags (Fisherbrand, catalog no. 15-182-501), and then flame-sealed to prevent escape. The bags were then submersed in media to stimulate mating and release of gametes. The medium formula consisted of the following reagents: HEPES buffering agent, L-Glutamine, Antibiotic/Antimycotic solution, and glucose.^8,16–18^ Nalgene wide mouth bottles containing media and cestodes were then placed into a shaking incubator at 42°C on slow speed. To test previous reports of decreased hatching of selfed eggs, we also generated a single clutch of selfed eggs from one Myvatn Lake parasite using the same protocol described above. The bottles were inspected daily to look for escaped worms and/or eggs. We harvest the eggs if they were present and the cestodes were transferred into a new bottle with fresh media and replaced in the incubator. Egg production usually stopped by the 4-5th day.^8,16–18^

### Density Treatments

Lab-bred cestode eggs representing two families from each population were distributed into 24-well plates across a range of densities. Test tubes containing *S. solidus* eggs in RO water were gently inverted to create a uniform distribution of eggs in the solution. Initially, density treatments were made in 24 well plates by pipetting 100µl, 200µl, and 400µl into six wells each for one outcrossed Walby egg solution and one outcrossed Myvatn egg solution. For the remaining outcrossed clutches, 50µl, 100µl, 200µl, 300µl, 400µl, and 500µl of the egg solution was transferred into four wells each of a 24-well plate (Figure 1). The lowest density treatment for one Walby plate was 10µl instead of 50µl, but all other density plates followed the scheme above. In total, we made two egg hatching plates from each cestode population representing one plate per family. Since the selfed crosses produce few eggs, we only examined a single plate from the selfed Myvatn cross. Each plate consisted of six density treatments, with 4 replicate wells of each treatment. Wells were then filled with RO water until they were at equal volumes and placed in a dark incubator set to 18□ for three days. Once removed from the dark incubator, plates were stored at room temperature under full spectrum light to induce hatching.

**Figure 1.**
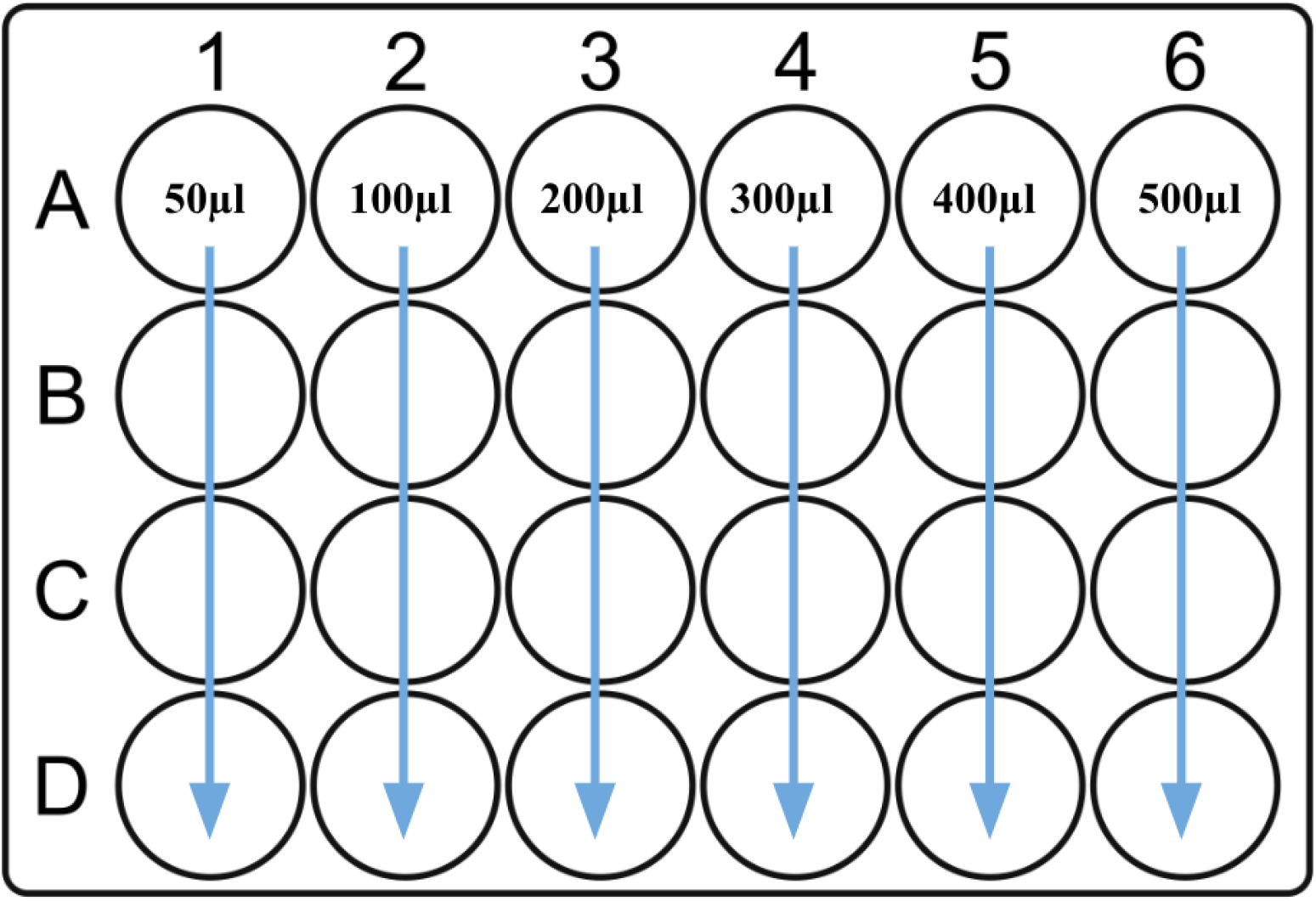
The plate layout used to assess if density impacts hatch rates of *S. solidus*. Volumes represent the total amount of thoroughly mixed egg stock added to each well.

### Measuring Egg Density and Hatching Rates

Plates were screened 2-3 times per week for hatching by scanning the water column for coracidia under a dissecting scope. Generally, hatching started ∼1 week after removal from the incubator and stopped 2-3 weeks later. Once the plate was finished hatching, an inverted scope was set to 4X and used to take a photo of each well. The location of each photo was decided by randomly selecting a location in the well when the microscope was not in focus. If the randomly selected location had less than 20 eggs in view, a new location was randomly chosen until the photo included at least 20 eggs. If all of the eggs were not in focus in a single photo, several photos were taken by first focusing on the eggs at the bottom of the water column and systematically moving to the top of the water column. If the photo was taken near the edge of the well, identical photos were taken with the brightness adjusted to ensure that all eggs were visible. The picture with the most eggs in focus was selected for counting. QuPath-0.5.1 (Bankhead, 2017) was used to manually count the number of hatched and unhatched eggs (Figure 2).

**Figure 2.**
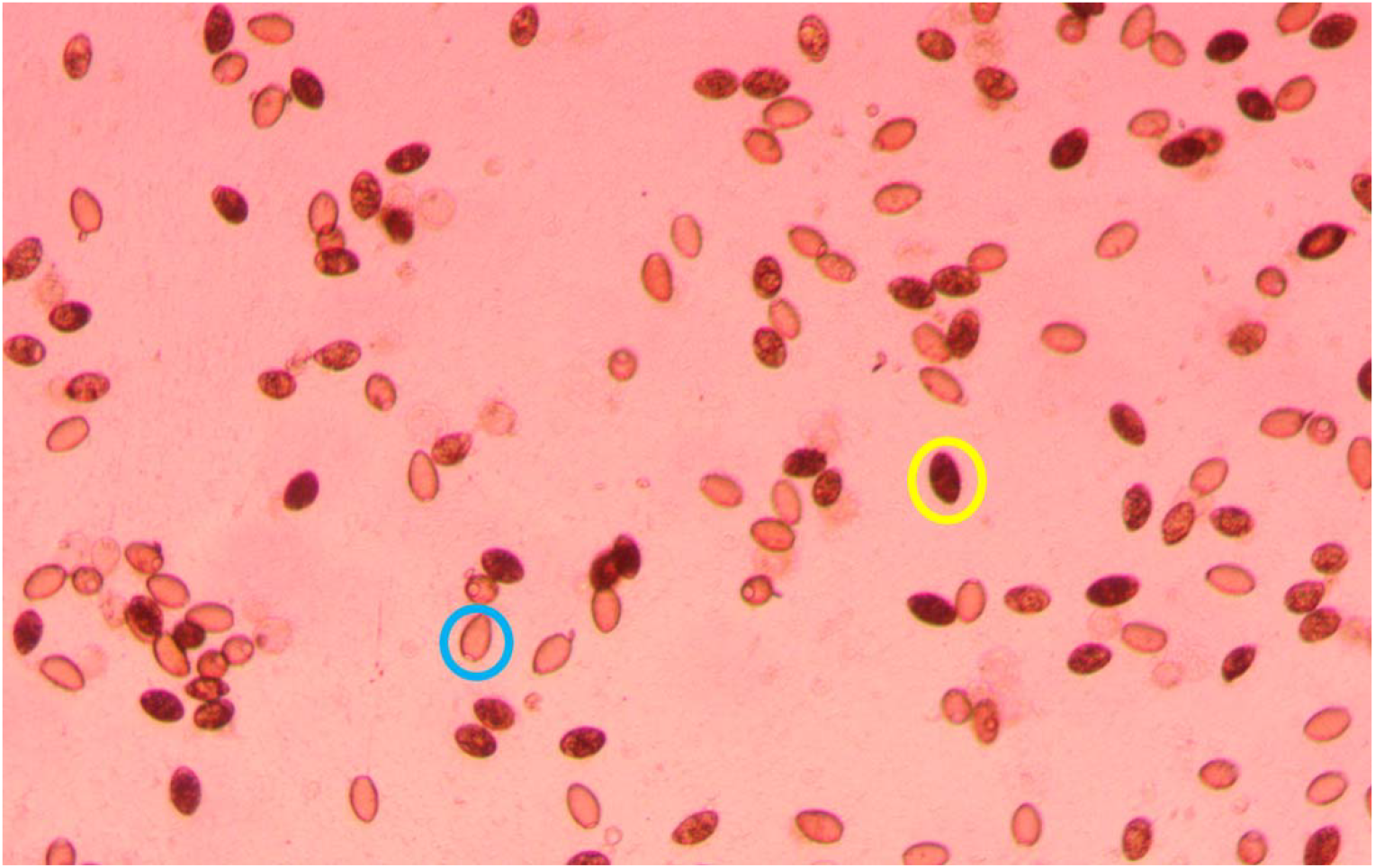
Photo of *S. solidus* eggs at 4x magnification, with an unhatched egg circled in yellow and a hatched egg circled in blue.

### Statistical analysis

All statistics were conducted in R v. 4.3.2.^19^ Counted eggs were log transformed so that the data was normally distributed. We fit a linear mixed effects model^20^ with p-values calculated using lmerTest v.3.1-3^21^ to determine if the volume of egg solution reliably predicted egg density with recorder as a random effect. We then used this model to predict the total number of eggs per well for each cross.^22^ A linear mixed effects model then was used to determine if the percentage of hatched eggs was influenced by the total number of eggs per well (i.e. egg density) or population, with cross and recorder as random effects. The analysis script and raw data are available at https://github.com/EmilyKerns/SSolidusEggDensity and Dryad. Photos of the eggs are available on Dryad.

## Results

We initially tested whether our method for aliquoting eggs led to a predictable increase in egg density. Of the six examined outbred crosses, a single Echo plate failed to hatch and was excluded from analyses. Additionally, while we did observe live coracidia while screening for hatched eggs from the single selfed cross, the hatching rate was so low that hatched eggs were difficult to find at 4x magnification. Thus, the selfed plate was also excluded from formal analysis. In total, we analyzed hatching rates from Walby (n = 2 crosses), Echo (n = 1 cross), and Myvatn (n = 2 crosses). There was variation across clutches in their starting densities which was not associated with population of origin. However, we observed a stable rate of increase of egg number with egg volume (t = 12.494, df = 0.0119, p<0.001; Figure 3A, Table 1). This demonstrates that volume was a suitable method for creating density treatments. We then used this model to predict the number of eggs in each density treatment for each cross (Figure 3B). The predicted number of eggs per well was used as an estimate of egg density in subsequent models.

**Table 1.** The results of each model used to assess hatching rates due to density both across all populations and within each population.

| Model | Fixed Effects | Estimate | Standard Error | df | t-value |
| --- | --- | --- | --- | --- | --- |
| Log(Total Number of Eggs) ~ Volume of Egg Solution + Cross + (1\|Recorder) | Intercept | 4.3266112 | 0.1290433 | 0.01190 | 33.528*** |
|  | Egg Solution Volume | 0.0038001 | 0.0003042 | 0.01190 | 12.494*** |
|  | CrossMyvatn3_1A | -0.7233042 | 0.1323678 | 0.01190 | -5.464*** |
|  | CrossMyvatn3_1C | -1.2401898 | 0.1447622 | 0.01190 | -8.567*** |
|  | CrossWalby_23_2 | 0.3706191 | 0.1447764 | 0.01190 | 2.560* |
|  | CrossWalbyBulk4 | -0.8863819 | 0.1590529 | 0.01190 | -5.573*** |
| Proportion Hatched ~<br>Density * Population<br>+ (1\|Cross) +<br>(1\|Recorder) | Intercept | 53.140206 | 14.168525 | 1.844381 | 3.751 |
|  | Density | -0.005158 | 0.010816 | 116.329217 | -0.477 |
|  | Myvatn | 22.004714 | 17.811145 | 2.116119 | 1.235 |
|  | Walby | -21.118892 | 17.138612 | 1.664919 | -1.232 |
|  | Density*Myvatn | -0.008165 | 0.024009 | 116.526355 | -0.340 |
|  | Density*Walby | -0.002827 | 0.013143 | 117.459670 | -0.215 |

\*\*\*p<0.0001, \*\*p<0.001, \*p<0.05

**Figure 3.**
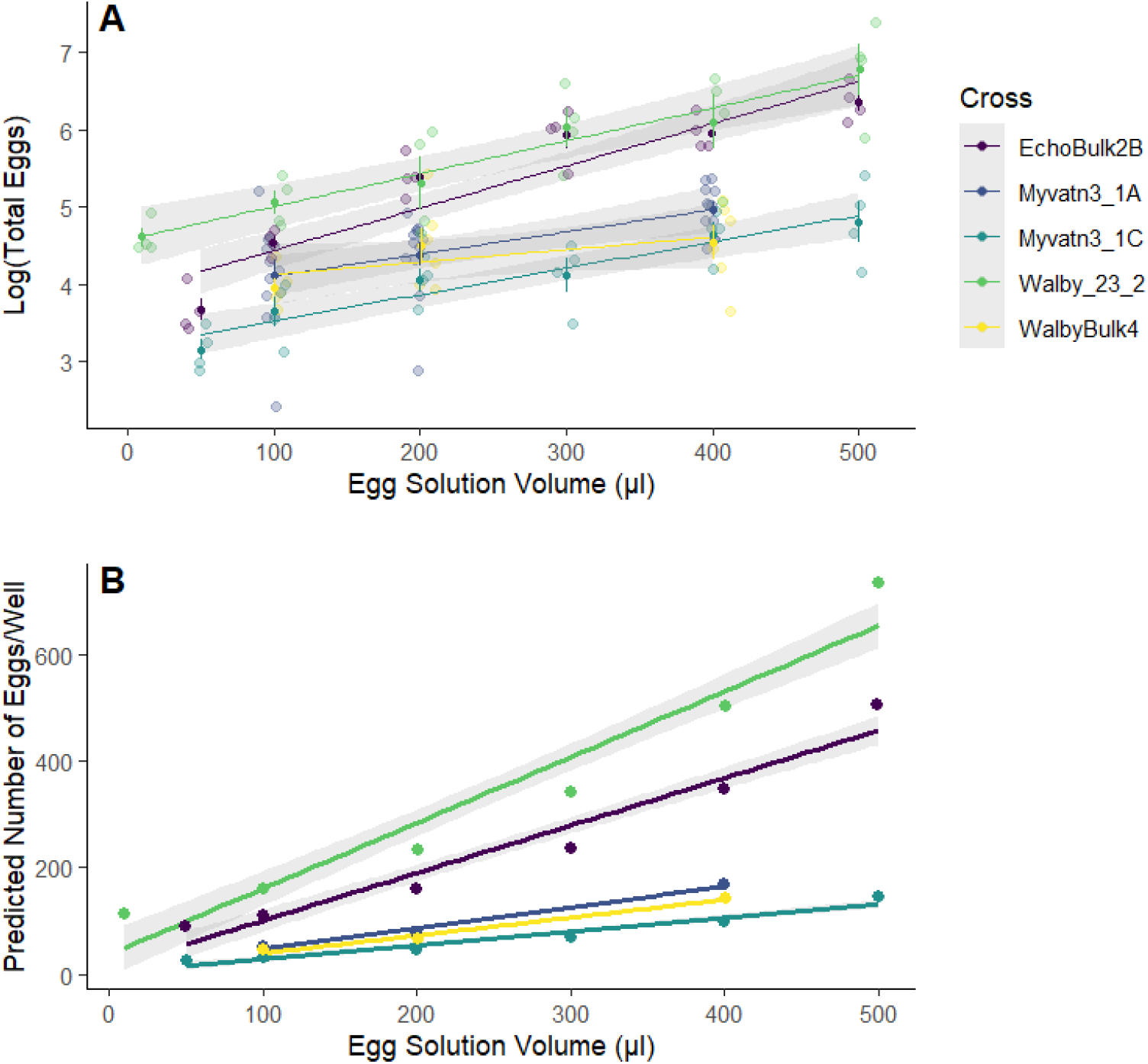
Density treatments based on egg solution volume. A) There is a significant positive association between the log transformed total number of eggs counted in each photo to the volume of egg solution used to create density treatments. B) The total number of eggs in each well was calculated using as model assessing the number of eggs per photo as a funciton of egg solution volume.

We next considered whether there was variation in hatching rates based on egg density, population of origin, and their interaction. A significant interaction term without a main effect of density would indicate that density-dependent hatching may be a derived trait in some populations. There was no effect of egg density on the proportion of hatched eggs or an interaction between populations. Additionally, while hatching rates were not significantly different between each of the populations, Myvatn tended to have reliably higher hatching rates than the other two populations (p > 0.05, Table 1, Figure 4). Although we did not formally measure whether egg hatching varied spatially within each well, we can anecdotally report that hatching did not seem to be clustered.

**Figure 4.**
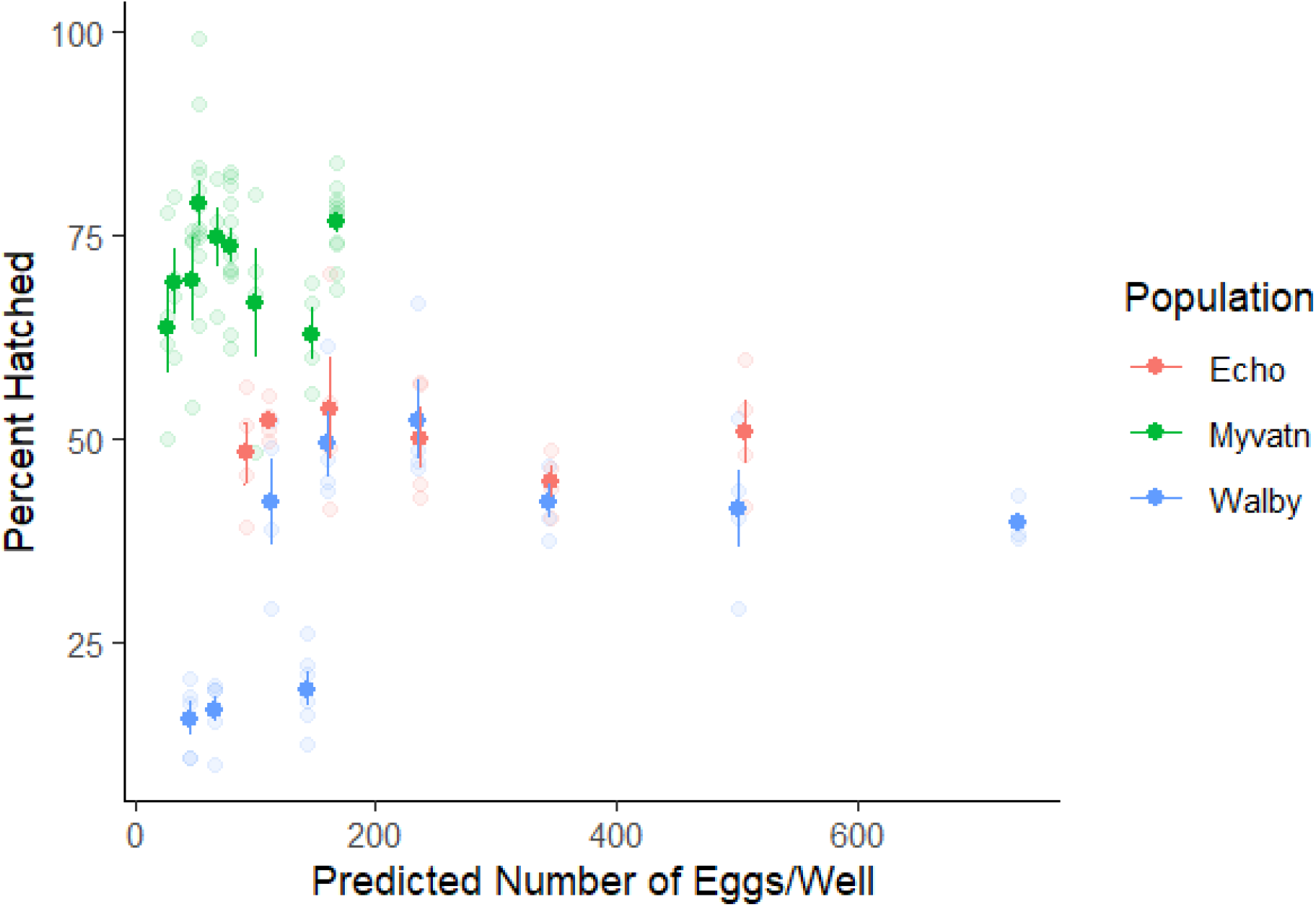
The relationship between egg density and hatching rates. There is no relationship between the predicted density of eggs per well and the proportion of hatched eggs.

## Discussion

Previous work has shown that selfing in the cestode *S. solidus* greatly reduces the hatching rate of progeny, suggesting that outcrossing should be preferred. However, previous studies have also shown that outcrossing only occurs when parasites encounter similarly sized mates.^23^ This is presumably because of fitness tradeoffs associated with anisogamy, where a larger cestode has to invest disproportionately more energy into the production of large eggs that can be cheaply fertilized by the sperm of the smaller mate. Integrating these results led us to hypothesize that selection may act to coordinate the timing of hatching, thereby increasing the probability of parasites encountering similarly aged (and thus similarly sized) mates. We also hypothesized that if egg hatching is accompanied by a signal that can induce the hatching of other eggs, then hatching rates may increase at high egg densities. This would be similar to density-dependent signaling observed in other organisms (i.e., QS). To our knowledge, apart from the effects of outcrossing versus selfing, no experiments have tested specific variables that may be associated with *S. solidus* egg hatching.

Although we were able to reliably generate and distribute *S. solidus* eggs across a range of densities, and also induce their hatching in our lab, we did not find evidence for density-dependent hatching. This negative result held both within and across three populations encompassing a broad geographic range. However, there was high variation in hatching rates between crosses within the same population and, while not statistically significant, variation between populations (Figure 4). Although our study was small and only assessed hatch rates in 3 outcrossed populations (≤ 2 families/population), these results suggest that hatch rate warrants further comparative study in this system.

Our original hypothesis was that synchronized egg hatching via QS might have evolved in *S. solidus* to increase rates of outcrossing. However, there are several lines of evidence that synchronized hatching and infection could be deleterious to this parasite. Previous studies found negative effects of crowding on *S. solidus* growth in both copepods^12^ and threespine stickleback^24^, indicating that coinfections decrease *S. solidus* fecundity by limiting growth across multiple intermediate hosts. Additionally, some populations of stickleback evolved the ability to encyst *S. solidus* in fibrotic tissue, thus preventing the cestode from growing and establishing an infection.^5,16^ Coinfection with multiple genotypes of *S. solidus* induces a higher fibrotic response than a single infection^25^, increasing the risk of parasite mortality. Therefore, costs of coinfection may outweigh the benefits of increased outcrossing, which would limit the potential for QS-based adaptation.

Despite the costs, QS-based hatching may still be selected for if the advantages of outcrossing outweigh the risks of coinfections. In agreement with previous work^13^, we observed a fitness deficit associated with selfing. The one selfed egg clutch that we examined contained relatively few eggs and did not produce sufficient hatch rates to be included in the study.

Stickleback infection rates vary drastically from lake-to-lake, with some populations having almost no infections.^16^ If transmission to the terminal host is rare, such as in lakes with low stickleback infection rates, coordinated hatching that leads to coinfection may still be advantageous to limit inbreeding. However, it is notable that *S. solidus* progeny produced after two consecutive generations of selfing had higher hatch rates than first-generation selfed progeny.^13^ The authors of this study speculated that this increase might result from purging of deleterious alleles, and that most lethal alleles could be purged from selfed lines in as few as 10 generations. Therefore, the high cost of selfing may reduce over time, even in extremely isolated populations of *S. solidus*. Future experiments on hatching rates and QS would benefit from explicitly choosing parasite populations that display low levels of heterozygosity, suggestive of long-term selfing. These are most likely to occur in geographically isolated lakes with relatively low infection rates.

Given that most of the data associated with QS come from bacterial studies, one explanation might be that this phenomenon is taxonomically biased. However, the mechanistic features of QS are highly similar to eukaryotic processes such as density-dependent autocrine signaling among cells within an organism.^26^ Indeed, numerous eukaryotes evolved to recognize, respond to, and manipulate the QS signals of bacteria.^27^ There is also strong evidence for intraorganismal signalling and QS in eukaryotes. The parasite *Trypanosoma brucei*, which causes African sleeping sickness, uses QS to vary developmental outcomes and increase transmission rates (Matthews, 2019). Similarly, the fungal pathogen *Cryptococcus neoformans*, the most common cause of fungal meningitis in humans, evolved a QS mechanism that mediates several pathways associated with its virulence.^28^ Although there may indeed be taxonomic variation in the presence and use of QS, specifically in the mechanisms and outcomes of signaling, we suggest there remains ample opportunity to connect the originally bacterial and highly molecular focus of QS to the broad ecological literature on positive density-dependent phenomena.^29,30^

## Conclusions

Although bacteria commonly exhibit QS, it is less commonly tested for in eukaryotes. We investigated the presence of QS in a eukaryotic parasite *S. solidus* using eggs from different worm populations and from different reproductive techniques (outcrossing versus self propagation). We found no evidence of QS between different concentrations of *S. solidus* eggs, but we did observe variation in overall hatching success between both reproductive techniques and some variation between populations. However, future studies of QS in *S. solidus* would be more robust by including more crosses and populations in the analysis and may unveil the evolutionary mechanisms that either prevent or facilitate adaptive QS.

## Acknowledgements

We would like to thank all members of the Weber Lab who assisted in copepod and *S. solidus* rearing. These *S. solidus* populations originate from the traditional and current lands of the Dena’ina and Coast Salish people. We thank them for their continued land stewardship and are grateful for the support of the local community. This work was funded by the Department of Integrative Biology at UW-Madison and National Institute of General Medical Sciences 5R35GM142891-02.

